# Peripherally derived LGI1-reactive monoclonal antibodies cause epileptic seizures *in vivo*

**DOI:** 10.1101/2023.10.11.561725

**Authors:** Manoj Upadhya, Toni Kirmann, Max Wilson, Christian M Simon, Divya Dhangar, Christian Geis, Robyn Williams, Gavin Woodhall, Stefan Hallermann, Sarosh R Irani, Sukhvir K Wright

## Abstract

One striking clinical hallmark in patients with autoantibodies to leucine-rich glioma inactivated 1 (LGI1) is the very frequent focal seizure semiologies, including faciobrachial dystonic seizures (FBDS), in addition to the amnesia. Polyclonal serum IgGs have successfully modelled the cognitive changes *in vivo* but not seizures. Hence, it remains unclear whether LGI1-autoantibodies are sufficient to cause seizures.

We tested this with the molecularly precise monoclonal antibodies directed against LGI1 (LGI1-mAbs), derived from patient circulating B cells. These were directed towards both major domains of LGI1, LRR (n=5) and EPTP (n=5) and infused intracerebroventricularly over 7 days into juvenile male Wistar rats using osmotic pumps. Continuous wireless EEG was recorded from a depth electrode placed in hippocampal CA3 plus behavioural tests for memory and hyperexcitability were performed. Following infusion completion (Day 9), post-mortem brain slices were studied using electrophysiology and immunostaining.

By comparison to control-mAb injected rats (n=6), video-EEG analysis over 9 days revealed convulsive and non-convulsive seizure activity in rats infused with LGI1-mAbs, with a significant number of ictal events (245±83 vs. 7.8±7.8 in controls; p=0.002). Memory was not impaired in the novel object recognition test. Local field potential recordings from postmortem brain slices showed spontaneous ictal-like spike activity in the CA3 region (p=0.03). The LGI1-mAbs bound most strongly in the hippocampal CA3 region and induced a significant reduction in Kv1.1 cluster number in this subfield (6 controls; 7 LGI1-mAbs; p=0.01)

Peripherally-derived human LGI1-mAbs infused into rodent CSF provide strong evidence of direct *in vivo* epileptogenesis with molecular correlations. These findings fulfill criteria for LGI1-antibodies in seizure causation.

## Introduction

Autoantibodies to leucine-rich glioma inactivated 1 (LGI1) are identified mainly in older males with a variety of distinctive and frequent focal seizure semiologies, including faciobrachial dystonic seizures (FBDS), piloerection and thermal seizures. In addition, many of these patients develop profound amnesia^1,2^. FBDS are characterised by brief and numerous seizures characterised predominantly by contractions of the arm and ipsilateral face ^3, 4^. If unrecognised and untreated, FBDS can evolve into a limbic encephalitis (LE) with temporal lobe and tonic-clonic seizures^4,5^, whereas, early treatment of FBDS with immunotherapies typically dramatically reduces seizure frequencies and can prevent the development of LE, thereby avoiding additional disability ^5, 6^.

Medial temporal lobe changes on MRI are a common finding in patients with LGI1-antibodies, and the subsequent atrophy predominantly affects the CA3 region ^7, 8^. Ictal EEG changes have been inconsistently reported, however, the most commonly recognised patterns are generalised attenuation, electrodecremental changes ^3, 9, 10^, simultaneous frontal EEG and EMG unilateral slow and infraslow wave preceding the contralateral tonic-dystonic seizure^11^, as well as frequent *subclinical* temporal lobe seizures ^3, 11, 12^.

Polyclonal patient serum-derived IgGs and LGI1-reactive monoclonal antibodies cause neuronal hyperexcitability and even epileptiform activity *in vitro*, with AMPAR and/or Kv1.1 downregulation postulated as the most likely underlying molecular mechanisms^13-15^. However, despite seizures being a key clinical hallmark of this disease, animal models using LGI1-antibodies have failed to recapitulate this pathognomonic feature *in vivo* ^16, 17^. This observation calls into question the direct epileptogenicity of LGI1-antibodies, particularly as the use of polyclonal human serum means that non-LGI1 reactivities may mediate observed outcomes.

Here, we use human-derived monoclonal antibodies with specific reactivities to either the leucine-rich repeat (LRR) or the epitempin repeat (EPTP) domain of LGI1^18^, to produce a passive transfer rodent model with spontaneous epileptic seizures and video-EEG recorded ictal changes.

## Materials and methods

### Antibody preparation

Two LGI1-specific mAbs, targeting either the LRR or EPTP domains were produced, as described previously ^18^.

### In vivo experiments

#### Animals

Sixteen postnatal day 21 (P21) male Wistar rats, weighing 52-60 g were used for experiments described. Animals were housed in temperature- and humidity-controlled conditions with a 12 h/12 h light/dark cycle and allowed free access to food and water. All procedures were compliant with current UK Home Office guidelines as required by the Home Office Animals (Scientific Procedures) Act 1986 and carried out under the authority and procedural approval of a UK Home Office approved project license and in line with ARRIVE guidelines. The Aston Bioethics Committee, University of Aston, Birmingham, UK granted local ethical approval for the study. Full experimental timeline shown in Figure 1A.

**Figure 1.**
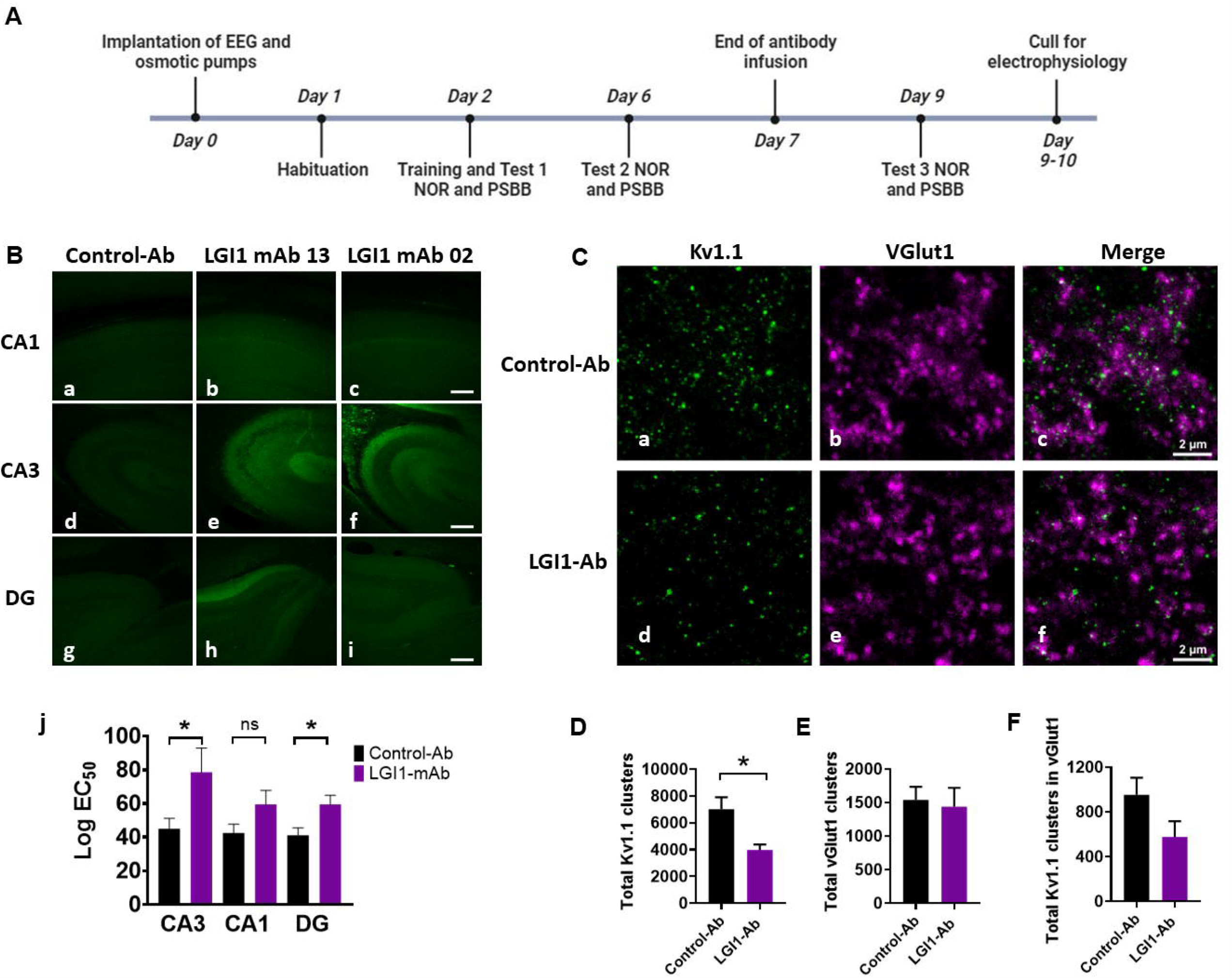
Patient derived monoclonal LGI1 antibodies bind most strongly to CA3 region of the hippocampus and cause a decrease of total synaptic clusters of Kv1.1 following 7-day intracerebroventricular infusion. (**A**). Experimental timeline. (**B**) (a-i) Representative confocal images of hippocampus (CA1 (a-c), CA3 (d-f) and DG (g-i)) from sagittal brain slice prepared after 7 days of infusion of Control-mAb (n=6; a, d, and f) and LGI1-mAbs (directed against epitopes EPTP n=3; (b, e, and h) and LRR n=4 (c, f, and i)) shows typical staining pattern with secondary anti-human IgG (green). Scale bar = 250µm. (Aj) LGI1 monoclonal antibody treated brain slices (17 slices) median fluorescent intensity log EC50 ratios compared to control-Ab (21 brain slices); *p<0.05, Mann-Whitney). (**C**) (a-f) Example images of STED microscopy of a section from the CA3 region stained for Kv1.1 (left) and VGlut1 (middle) as well as the merge (right) fromm brains treated with control-Abs (top) or LGI1-Ab (bottom). (**D**) Quantification of density of total Kv1.1 clusters in pooled analysis of CA3 region in animals treated with Control or LGI monoclonal antibodies. (**E**) Quantification of density of total vGlut1 clusters in pooled analysis of CA3 region in animals treated with Control or LGI monoclonal antibodies. (**F**) Quantification of total KV1.1 clusters colocalizing with vGlut1. C-E (n= 5 images of CA3 region for each brain sample: 6 controls; 7 LGI1 mAbs (4 LRR specific mAb2 and 3 EPTP specific mAb13). Data are presented as mean ± SEM. \**P* < 0.05, \*\**P* < 0.01, \*\*\**P* < 0.001.

### EEG implantation and osmotic pump surgery

The rats were implanted, as previously described, with subcutaneous transmitters for *in vivo* EEG recordings (A3028B-DD subcutaneous transmitters, 90-mm leads, OpenSource Instruments (OSI)) via unilateral depth electrode (W-Electrode (SCE-W), OSI) in left hippocampus (CA3, 3.5 mm lateral, 3.8 mm caudal, depth 2.3 mm), and subcutaneous osmotic pumps (model 1007D, Azlet) (volume 100 µl, flow rate 0.5 µl/hr, duration 7 days, primed overnight) attached to bilateral cannulae (328OPD-3.0/SPC; PlasticsOne), implanted into the lateral ventricles (±1.5 mm lateral and 0.6 mm caudal) ^19-21^. A reference EEG electrode was implanted on the contralateral skull surface (3.5 mm lateral, 3.8 mm caudal); the cannula and skull electrodes were secured with dental cement ^19, 20^.

### EEG data analysis

EEG data (wide band pass 0.2–160◻Hz sampled at 512 samples per second) was collected and recorded using Neuroarchiver software (OSI) from wireless transmitters implanted in freely moving animals placed in a custom-built Faraday cage with aerial (OSI) as previously described ^19,20,21^. For automated ictal event detection, video-EEG matching was used to identify ictal EEG events. The Event Classifier (OSI) was used to classify segments (1 second) of EEG according to program metrics (power, coastline, intermittency, coherence, asymmetry, spikiness) creating clusters of similar events when plotted. This generated a library of ictal events that allowed fast identification of abnormal EEG events by automated comparison (http://www.opensourceinstruments.com/Electronics/A3018/Seizure_Detection.html). Powerband analysis was carried out using a custom-designed macro. Statistical analysis was conducted using Graphpad Prism 8 (GraphPad Software Inc.).

### Behavioural testing

#### Novel Object Recognition (NOR)

The novel object recognition test is used to test cognitive performance in autoimmune encephalitis models ^17, 22^. Animals were habituated to the test arena 24 hours before training and tests. The time interacting with the objects was measured and the NOR index (time interacted with novel object/ total time of interaction with both the objects) calculated. Three tests were carried out at regular intervals during the 9-day recording period. All the observations were carried out by the experimenter blind to the treatments and analysed using the Ethovision software. The frequency of rats to approach the object, time spent with the objects was measured for NOR index, and the distance travelled, and velocity was measured as a measure of anxiety score. A repeated-measures 2-way ANOVA with Bonferroni multiple comparison test and non-parametric Mann-Whitney T-test was used to assess differences between NOR index, distance travelled and velocity for different cohorts. P<0.05 was considered as significant.

#### Post-Seizure Behavioural Battery (PSBB)

The Post-Seizure Behavioural Battery (PSBB) was performed as previously described to assess hyperexcitability and other behavioural indices suggestive of epilepsy ^20, 23^. Two simple and non-stressful tasks, “touch” and “pick-up” tasks constituted the PSBB and were performed at three time-points. PSBB scores were calculated by taking the product of the task scores (‘touch x pickup’). A repeated-measures 2-way ANOVA with Bonferroni multiple comparison test was used to assess differences between PSBB scores for different cohorts.

## In vitro experiments

### Local field potential recordings

#### Local field potential (LFP) recordings

LFP recordings were performed and analysed as previously described ^19, 20^. Briefly, on day 9 immediately after the behavioral experiments, rats were anaesthetized using isoflurane and following the loss of consciousness, pentobarbital (60 mg/kg, SC) and xylazine (10 mg/kg, IM) injected. Transcardial perfusion was then performed using ice-cold artificial cerebrospinal fluid (aCSF). Animals were decapitated, and the brain was removed. Brain slices were prepared using a vibratome (Campden Instruments, UK) at 450μm for LFP recording and 350μm for fluorescence intensity measurements. LFP recordings were assessed for spontaneous epileptiform activity using Spike2 software (CED). Root mean square (RMS) amplitude of each recording was calculated, and epileptiform activity was classified as an event (“spike”) when it displayed an amplitude greater than four-fold the RMS amplitude. Statistical analysis was conducted using Graphpad Prism 8 (GraphPad Software Inc), statistical tests used to compare groups (Mann-Whitney) were one-tailed.

### Immunofluorescence, immunohistochemistry and image analysis

The immunofluorescence study was performed as previously reported ^19, 20^. Further detailed methods available in Supplementary information.

### Data availability

Data are available on request.

## Results

### LGI1 antibodies bind to rat hippocampus and cause reduction of Kv1.1 clusters

Seven-day osmotic pumps delivered the LGI1-mAbs (n=10; five animals with LRR and five with EPTP specific mAbs) and control mAbs (n=6) into the lateral cerebral ventricles of P21Wistar rats.

After interventricular infusion, both LGI1-mAbs (LRR and EPTP specific) bound to rodent hippocampus (Figure 1B), confirming previous studies in mice ^18^. By comparison to control, non-brain-reactive mAbs, the fluorescence intensities of bound LGI1-mAbs were highest in the CA3 and dentate gyrus. In the CA3 region, super-resolution light microscopy using stimulated emission depletion (STED) of 20-µm-thick cryosections revealed that the number of Kv1.1 clusters was significantly lower in the LGI1-mAb infused group (pooled analysis of LRR and EPTP) as compared to controls (Figure 1C), and non-significantly trended to those which co-localised with the glutamatergic synapse marker vGlut1 (Figure 1D, E).

### LGI1-mAbs cause clinical and subclinical spontaneous epileptic seizures *in vivo*

To investigate potential epileptogenic effects, brain activity was recorded continuously during LGI1-mAb infusion using wireless EEG transmitters from a depth electrode placed in the CA3 region of hippocampus. One rodent infused with EPTP specific LGI1-mAb was culled early due to excess lacrimation, and in one rodent infused with LRR specific mAb, the EEG recording was suboptimal for analysis but the animal data were still included for behavioural testing and post-mortem immunohistochemistry. Overall, the recorded epileptic and behavioural activity included convulsive, non-convulsive and inter-ictal events (Figure 2A i-iii, Suppl. videos 1,2). Using automated ictal event detection ^19, 20, 21^, we observed a significantly higher total number of ictal events in the LGI1-mAb infused animals compared to controls during the 9 days of EEG recording period (Figure 2B). EEG coastline was higher in LGI1-mAb infused animals, due to higher prevalence of relatively high-amplitude epileptiform events (Figure 2C); this cohort also showed higher power in all EEG frequency bands (Figure 2D). Following completion of the antibody infusion, 4 LGI1 and 4 control mAb infused animals were terminally anaesthetized and brains extracted for acute *in vitro* brain slice recording. Local field potentials were recorded in CA3 and CA1 of the hippocampus (Figure 2E, F), and showed spontaneous epileptiform activity in the brain slices prepared from LGI1-mAb infused animals.

**Figure 2.**
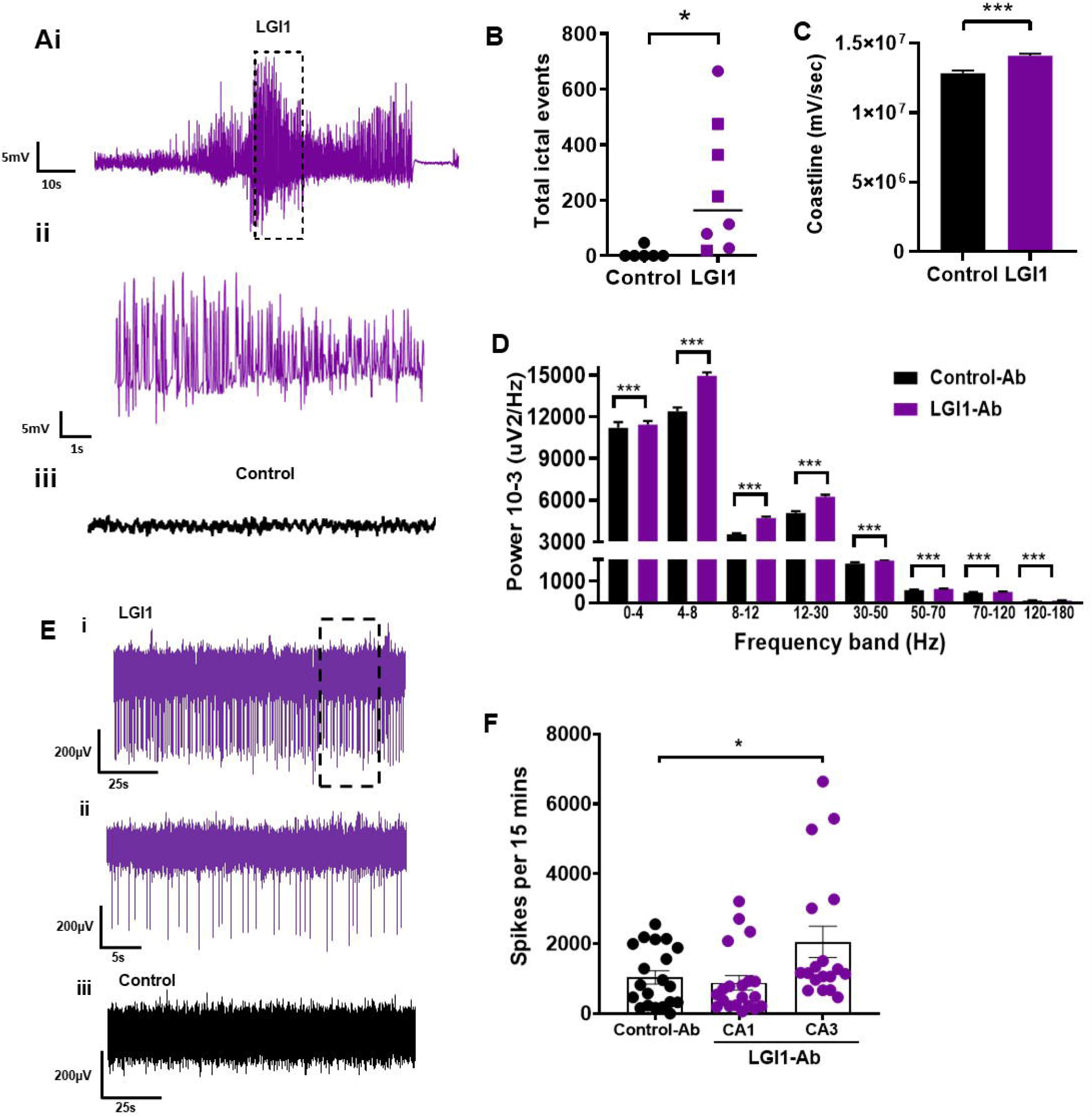
Intracerebroventricular infusion (7 days) of patient derived monoclonal LGI1 antibodies induces seizures and epileptiform activity in vivo and in vitro. **(A)** Example EEG recording from rodent using wireless EEG transmitter during intracerebroventricular infusion of LGI-mAb directed towards LRR (Suppl Video1) (i), highlighted/hatched EEG area expanded in (ii); control mAb infused rodent EEG shown for comparison in (iii). **(B)** Total number of 1-second ictal events recorded during 7-day intracerebroventricular infusion of LGI-mAbs compared (n=8; 4 LRR epitope and 4 EPTP) to control-Ab infused animals (n=6). Squares represent rodents infused with LRR epitope specific LGI1-mAb, circles represent EPTP epitope specific LGI1-mAb. Data are presented as mean ± SEM. (Mann-Whitney, \**P* < 0.05). **(C)** Averaged hourly EEG coastline length for LGI1-mAb infused rodents (n=8; 4 LRR epitope and 4 EPTP) compared to controls (n=6) (Mann-Whitney, \*\*\**P* < 0.001). **(D)** Hourly EEG power averages over 7-day infusion and recording period of LGI1-mAbs versus controls (Mann-Whitney, \*\*\**P* < 0.001); rodent numbers as in (B,C). **(E)** Example traces of local field potential brain slice recordings from LGI1-Ab (i-ii) and control-Ab (iii) infused rats for 7 days and culled on day 9 from the CA3 region. Scale bar 200◻µV vs. 25◻s (i), 200◻µV vs. 5◻s (ii) and 200◻µV vs. 25◻s (iii). **(F)** The number of spikes (interictal events) per 15◻min in the brain slices infused with LGI1-Ab compared to control-Ab infused slices (Control-Ab (n◻=◻10 brain slices) vs. LGI1-Ab (n◻=◻20 brain slices for CA1 and 18 brain slices for CA3); *p◻=◻0.0314, Mann–Whitney). Data are presented as mean ± SEM.

### LGI1-mAbs cause ictal hyperexcitable behavior without effect on cognition

The post-seizure behavioural battery (PSBB) was used to monitor behavioural changes (hyperexcitability and aggression) consistent with the development of spontaneous recurrent seizures ^20, 23^. Animals were tested three times over 9 days using touch and pick-up tests, with a score > 10 indicating significant behavioural change. The LGI1-mAb infused animals scored significantly highly at all three time-points compared to the control-Ab infused animals where the scores remained consistently low (Figure 3A).There were no differences in the NOR index (calculated by dividing the time spent with novel object to the entire duration of time spent with both the objects);, locomotion, and distance travelled and velocity, during NOR across three timepoints (day 1, day 5 and day 9; Figure 3B-D).

**Figure 3.**
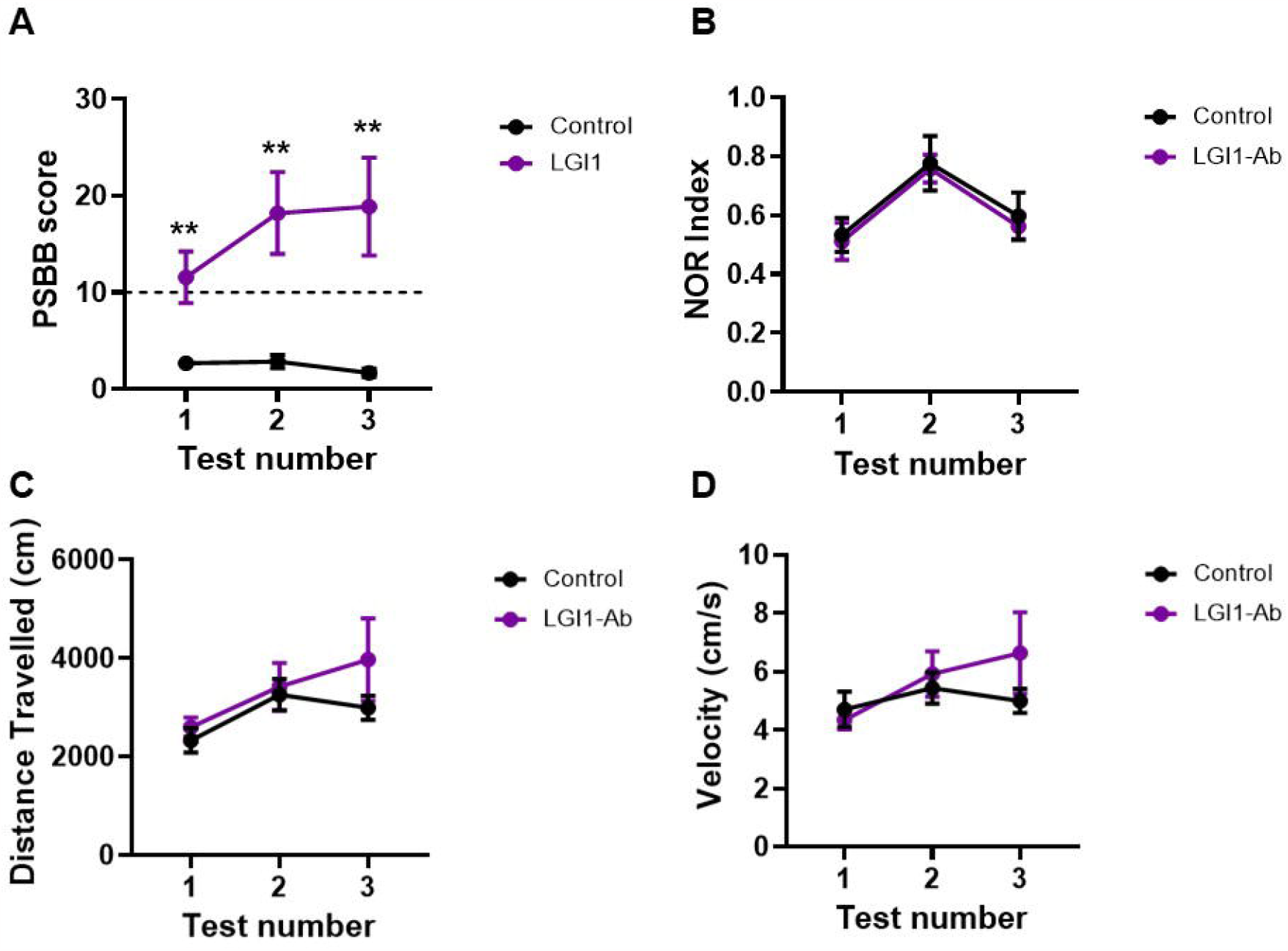
Intracerebroventricular infusion (7 days) of patient derived monoclonal LGI1 antibodies causes hyperexcitability but has no effect on memory in behavioural tests. **(A)** Post-seizure behavioural battery scores in rodents over 7-day intracerebroventricular infusion of LGI-mAbs (n=9; 5 LRR epitope and 4 EPTP) and control-Ab (n=6) (one-way ANOVA, \*\**P* < 0.01); dotted line represents a score of 10, over this is considered suggestive of hyperexcitable behaviour. **(B)** Novel object recognition (NOR) index calculated at three timepoints of LGI1-mAbs (n=7; 4 LRR epitope and 3 EPTP) and control-mAb (n=6) infused rats. **(C)** Distance travelled by rats and **(D)** velocity measured during NOR tests in rats infused with control-mAb (n=6) and LGI1-mAbs (n=7; 4 LRR epitope and 3 EPTP) infused rats. Data are presented as mean ± SEM.

## Discussion

LGI1-antibody encephalitis patients exhibit focal and tonic-clonic seizures as well as cognitive impairment. Published animal models were based on application of serum-IgGs to successfully model the cognitive changes *in vivo*, but seizures were not seen ^16, 17^. Here, we used highly specific patient-derived LGI1-mAbs, which lack the potential additional reactivities potentially present in serum. After infusion into the CSF of juvenile Wistar rats, these LGI1-mAbs led to seizures *in vivo*. Similar to patient phenotypes, we recorded clinical and subclinical seizures, with ictal and interictal EEG changes. These findings support the direct epileptogenicity of LGI1 antibodies and allow them to fulfill Wietebsky’s postulates of pathogenicity ^24^.

LGI1 is a secreted neuronal protein that interacts with the ADAM22 and ADAM23 transmembrane metalloproteases forming a trans-synaptic protein complex that facilitates excitatory synaptic transmission ^25-27^. LGI1 consists of an N-terminal LRR domain and a C-terminal EPTP domain. The LRR-directed monoclonal antibody (mAb) used in this study binds the ADAM22/23-docked LGI1 complex *in vitro*, causing internalisation and hence disruption of the membrane stabilising trans-synaptic complex ^18^. This pathomechanism is similar to that demonstrated by a CSF-derived LGI1-specific mAb that also caused epileptiform changes *in vitro*^14^. *In vitro* primary neuronal cultures also incubated for 7 days with LRR and EPTP specific mAbs showed that LRR-directed mAbs could directly affect neuronal excitability, but this effect was less pronounced in EPTP-specific mAb exposed neurons ^28^. In contrast to the action of LRR mAbs *in vitro*, the EPTP mAbs are reported to exert their effects by *directly* inhibiting the docking of LGI1 to the ADAM proteins ^18^. In our study, EEG and LFP recordings have demonstrated increases in network and neuronal excitability in hippocampal CA3 *in vivo* and *in vitro* by both LRR and EPTP domain directed LGI1 autoantibodies ^14, 29^. Studies have shown that LRR- and EPTP-specific antibodies co-occur in patients with LGI1-antibody encephalitis but the numbers of recorded rats with either mAb subclass do not allow for comparison of epitope-specific clinical features ^14^. The goal of future studies is to analyse our *in vivo* EEG using computational modelling to evaluate different hypotheses pertaining to network hyperexcitability and seizures by action of epitope specific LGI1-antibodies and to differentiate epitope-specific effects ^19, 30, 31^.

EPTP specific mAbs are reported to have no effect on cognitive performance when injected into hippocampi of mice ^18^: this may explain why our pooled NOR results from both LRR and EPTP specific mAbs infused rats were unremarkable. Additionally, in contrast to previous *in vivo* passive transfer models, our antibody infusion time was much shorter: 7 days, compared to 14 days. Patients with LGI1-Ab encephalitis frequently develop seizures before the onset of memory disturbance ^32^ and this could explain our behavioural findings in this rodent model. Lastly, the genetic background of rodents can impact on seizure susceptibility ^16, 21, 33^; here juvenile Wistar rats were used as they have proven to be effective in modelling autoimmune-associated seizures and epilepsy *in vivo* ^19, 20^.

The goal of this study was to prove epileptogenicity of LGI1 mAbs *in vivo* using mAbs with known pathogenic effects *in vitro*. We also took this opportunity to generate some insights into identifying a pre- or post-synaptic mechanism for seizure generation. Previous longer-term infusion (14 days) *in vivo* studies using patient-derived LGI1-mAbs that target both the LRR- and EPTP domains did show a disruption in presynaptic and postsynaptic LGI1 signalling ^17^. This was due to an initial decrease of Kv1 levels (at 13 days) followed by AMPA receptor downregulation by 18 days post-infusion ^17^. We also demonstrated loss of Kv1.1 clusters in hippocampal CA3 (with a trend towards a reduction in glutamatergic synapses) but our infusion time was shorter (7 days), and longer-term timepoints were not analysed.

In summary, in this focused study, we have developed a passive transfer *in vivo* model of LGI1-mAb associated seizures which displays epileptiform activity in keeping with the human disease. The use of wireless EEG telemetry and continuous video-EEG allows for accurate tracking of behavioural and EEG changes. Future studies will focus on interrogating new and existing data on exact pathomechanisms for *in vivo* epileptogenicity and identification of possible novel epitope-specific treatment targets.

## Supporting information

Suppl video 1 convulsive

Suppl video 2 non-convulsive

## Abbreviations

AMPAR: α-amino-3-hydroxy-5-methyl-4-isoxazole propionic acid
CSF: cerebrospinal fluid
EPTP: epitempin repeat
FBDS: Faciobrachial dystonic seizures
IgG: Immunoglobulin
LE: Limbic encephalitis
LGI1: leucine-rich glioma inactivated 1
LRR: leucine-rich repeat
NOR: Novel object recognition
PSBB: Post seizure behavioral battery

