## Supplementary information for "Peripherally derived LGI1-reactive monoclonal antibodies cause epileptic seizures *in vivo*"

**Supplementary Data**

### Immunofluorescence, immunohistochemistry, and image analysis

Brain slices (350 µm) were briefly fixed in 4% paraformaldehyde (PFA) for 60 min. To determine LGI1 or control antibody binding, the sections were rinsed with phosphate-buffered saline (PBS) and then incubated in antihuman IgG Alexa-Fluor-488 at 1 in 1000 (Invitrogen, UK) overnight at 4 °C. Sections were washed and mounted with aqueous mounting medium containing DAPI (Vectashield). Images of hippocampal sections were taken on a Tandem Confocal Scanning SP5 II microscope (Leica Microsystems Ltd) using a (10×/0.30) dry objective lens. Fluorescence was excited with a 488 nm argon laser at 20% power (emission bandwidth 504nm-564nm). Confocal micrographs were acquired at 1024 pixels^2^ with actual area of 1.48mm^2^ and scanning speed 100 Hz.

Fluorescent intensity Log EC50 ratio of hippocampal sections was determined using 5 regions of interest (ROI) of the same size from CA1, CA3 and DG regions. The mean fluorescent intensity Log EC50 value was calculated by nonlin fit method using GraphPad Prism 8. All images were analysed through FIJI by converting them to grayscale (8-bit) before generating cumulative pixel intensity histograms for each ROI which were calculated using a customized macro programme. Statistical analysis performed was conducted in GraphPad Prism 8.

For immunohistochemical analysis of the rat brains, one hemisphere was perfused, fixated in 4% PFA, cryoprotected in 30% sucrose and embedded in optimal cutting temperature compound. They were then rapidly frozen in isopentane cooled with liquid nitrogen.

To assess the levels of presynaptic Kv1.1 potassium channel brain slices of 20 µm thickness were prepared using a cryotome. Excess OCT medium was removed by washing the sections once with PBS at room temperature for 5 min. The samples were then subjected to a 1 hour blocking step (block buffer: 1xPBS + 0.3% Tx-100 + 5% NGS + 5% NDS) at room temperature followed by an overnight incubation at 4°C with antibodies against Kv1.1 rabbit IgG (1:250, AB_2571788, Nittobo Medical), Bassoon mouse IgG (1:400, SAP7F407, Enzo Life Sciences) and vGluT1 guinea pig IgG (1:2500, 135304, Synaptic Systems). The following day, the sections underwent a PBS washing step, involving three rounds of washing, each lasting 5 minutes at room temperature. Subsequently, the samples were counterstained at room temperature for 1.5 hours using secondary antibodies, namely goat anti-rabbit StarRed, goat anti-mouse StarOrange, and goat anti-guinea pig Alexa Fluor 488. The sections were then incubated with a DAPI stain for 20 min at room temperature before mounting and sealing with nail polish in a 7:3 PBS:Glycerol solution.

For analysis of the total number of Kv1.1 signals and their co-localisation with vGluT1, we used an Expert Line Abberior microscope system with an Olympus (UPlanXApo) 60x oil objective with a numerical aperture of 1.42. The system contained 3 pulsed fluorescence excitation laser modules for wavelengths of 485, 561, and 640 nm, the corresponding filter cubes (GFP, Cy3, Cy5), and avalanche photodiode (APD) detectors. For STED imaging the system uses a pulsed high-power STED laser module with a wavelength of 775 nm (3 W laser power, repetition rate of 40 MHz). Each image was obtained with sequential line scanning by switching between the needed excitation lasers line-wise during each recording. Each image contained a three channel-recording for Kv1.1, the active zone protein Bassoon and the vesicular glutamate transporter vGluT1. All channels were imaged with a dwell time of 5 μs/pixel and a onefold line accumulation. Based on the secondary antibody signals, the excitation laser power for each channel was adjusted to prevent over- or underexposure effects. Excitation laser power for Kv1.1 (secondary antibody, STAR Red, laser module 640 nm) was set to 20% with a STED laser power of 20% and for Bassoon (secondary antibody, STAR Orange/CF568, laser module 561 nm) to 70% with a STED laser power of 20%. For vGluT1 (secondary antibody, Alexa Fluor 488, laser module 488 nm) only confocal images were acquired with an excitation laser power of 70%. The settings were kept constant throughout all image acquisitions.

For quantification, a custom-written macro of the software Fiji43 was used to generate (1) a mask based on a prior adjusted threshold for Kv1.1 to count Kv1.1 spots and (2) a mask for vGlut1 to quantify the average pixel intensity of Kv1.1 within the vGluT1 masks.

Supp Video 1

Convulsive seizure captured from rodent infused with LGI1 monoclonal antibodies directed towards the EPTP domain. The EEG trace accompanying the seizure is shown in Fig Ai.

Supp Video 2

Non-convulsive seizure captures from rodent infused with LGI1 monoclonal antibodies directed towards the LRR domain, behavioural arrest seen from 23 seconds. The EEG trace accompanying the seizure is shown below. Scale bar 5mV vs 60s.

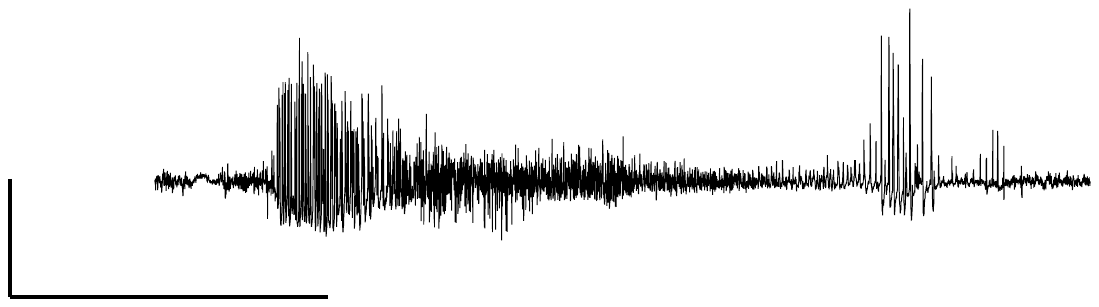
